# Monitoring the promoter activity of long noncoding RNAs and stem cell differentiation through knock-in of sgRNA flanked by tRNA in an intron

**DOI:** 10.1101/2021.01.08.425823

**Authors:** Yu-Ting Zhao, Yangming Wang

## Abstract

Mammalian genomes encode thousands of long noncoding RNAs (lncRNAs) that are often expressed in a tissue and cell specific manner. Therefore, a reporter that can faithfully reflect the expression or activity of lncRNAs can provide tools useful not only for uncovering the regulators of lncRNAs, but also for tracking cell fate and disease status. Here, we design a s<u>gR</u>NA precursor in an <u>i</u>n<u>t</u>ron (GRIT) strategy that can monitor the promoter activity of lncRNAs. We used this strategy to report the expression of Lncenc1 and Neat1 in mouse embryonic stem cells (ESCs). Furthermore, we show that GRIT may be used to track differentiation status of stem cells. We anticipate that GRIT will be applicable in dissecting regulatory mechanisms underlying the transcription of lncRNAs, tracking cell fate switch during differentiation or disease progression and integrating the promoter activity of various RNAs for synthetic biology applications.

---

The majority of mammalian genome is transcribed to RNA transcripts, of which only a very small percentage code for proteins^1^. As a result, thousands of RNAs that do not code for proteins are produced in cells, including microRNAs (miRNAs) and long noncoding RNAs (lncRNAs). These noncoding RNAs exert regulatory functions in various physiological and pathological conditions^2^. In addition, numerous noncoding RNAs are expressed in a tissue and cell specific manner^1^. Thus, a reporter that can faithfully reflect the expression or activity of noncoding RNAs can provide tools useful not only for uncovering the regulators of noncoding RNAs, but also for tracking cell fate and disease status. Previously we have designed a miRNA inducible CRISPR-Cas9 platform that can serve as a sensor for miRNA activities^3^. However, designing a reporter for long noncoding RNAs has not been easy due to its untranslatable nature and low expression level. Here, we design a s<u>gR</u>NA precursor in an <u>i</u>n<u>t</u>ron (GRIT) strategy that can monitor the promoter activity of lncRNAs (**Fig. 1a**). Furthermore, we show that GRIT can be used to track differentiation status of stem cells.

**Figure 1.**
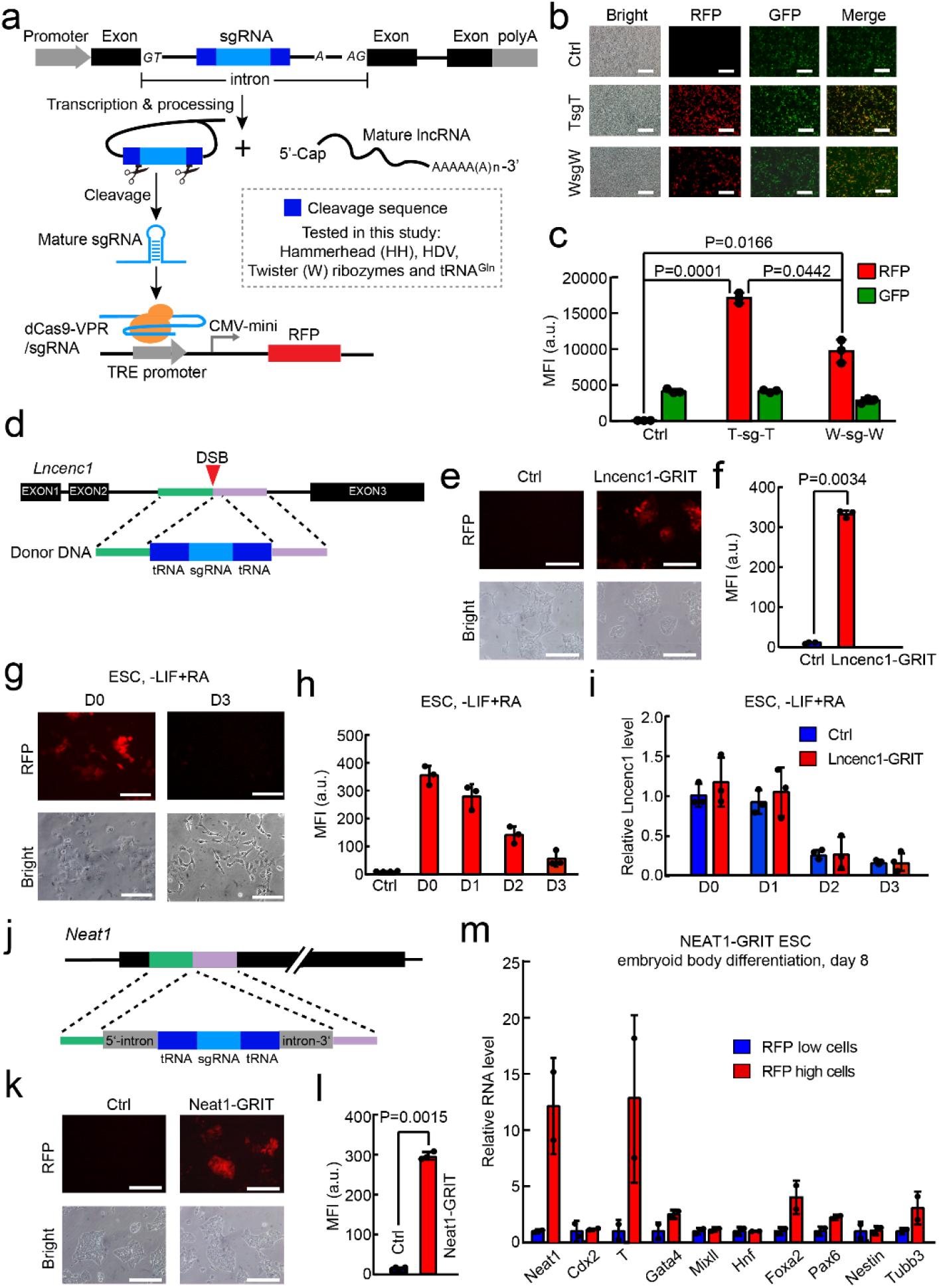
GRIT monitors the promoter activity of lncRNAs and stem cell differentiation. **a)** Schematic design of GRIT reporter system. GRIT cassette refers to pre-sgRNAs located in an intron. After the transcription of host gene, removal of flanking RNA cleavage sequences leads to the maturation of sgRNA which activates the downstream CRISPR-on reporter system. Hammerhead and HDV ribozyme sequences from Ref. 8; Twister ribozyme sequence from Ref. 9. **b)** Representative microscopy images showing RFP and GFP expression in HEK293T transfected with dCas9-VPR, TRE3G-RFP and GRIT-GFP plasmids. The schematic for the design of this experiment is shown in **S1a**. Scale bars: 200μm. The experiments were repeated three times independently with similar results. TsgT, tRNA-flanked gRNA. WsgW, Twister ribozyme-flanked gRNA. **c)** Quantification of mean RFP and GFP intensity of **b**. Shown are mean ± SD, n = 3 independent experiments. The p-value was calculated by one-way ANOVA with two-tailed Tukey’s multiple comparisons test. **d)** Schematic of GRIT knock-in strategy for *Lncenc1*. After the establishment of dCas9-VPR and TRE3G-RFP transgenic mouse ESCs, the TsgT element is knocked in the second intron of *Lncenc1* locus through CRISPR-Cas9-assisted homologous recombination. **e)** Representative images showing RFP expression in Lncenc1-GRIT ESCs. **f)** Mean RFP intensity of Lncenc1-GRIT ESCs and control ESCs which are dCas9-VPR and TRE3G-RFP transgenically integrated but without knock-in of Lncenc1-GRIT. Shown are mean ± SD, n = 3 independent experiments. The p-value was calculated using two-tailed unpaired Student’s t-test. **g)** Representative images showing RFP expression in undifferentiated and differentiated Lncenc1-GRIT ESCs. Scale bar, 200 μm; **h)** Quantification of mean RFP intensity during the continuous differentiation process of Lncenc1-GRIT mESCs. Shown are mean ± SD, n = 3 independent experiments. **i)** RT-qPCR analysis of Lncenc1 expression during differentiation process of Lncenc1-GRIT and control ESCs. Shown are mean ± SD, n = 3 independent experiments. **j)** Schematic of GRIT knock-in strategy for *Neat1* locus. An artificial intron based on the second intron of human *RPL18* gene containing GRIT elements was knocked into the *Neat1* locus. **k)** Representative images showing RFP expression in Neat1-GRIT cells. **l)** Mean RFP intensity of Neat1-GRIT ESCs. Shown are mean ± SD, n = 3 independent experiments. The p-value was calculated using two-tailed unpaired Student’s t-test. **m)** RT-qPCR analysis of various differentiation markers in RFP high and low cells from day 8 differentiating embryoid bodies of Neat1-GRIT ESCs. EB, embryoid body. Shown are mean ± SD, n =2 independent experiments.

The design of GRIT includes three key elements (**Fig. 1a**): dCas9-VPR expressed under the control of a constitutively active CAGGS promoter^3^, a RFP gene under the control of a tetracycline-inducible promoter^3^, and a transfer RNA^Gln^ (tRNA^Gln^)^4^-flanked sgRNA that is integrated in an endogenous noncoding RNA locus through homologous recombination. To minimize the impact of tRNA-sgRNA knock-in on lncRNAs, we choose genome region that will be expressed as an intron to knock in sgRNAs. In addition, for lncRNA gene without an intron, an artificially designed mini intron containing tRNA-sgRNA fusion sequence is knocked in. The tRNA flanking design was chosen based on our observation that tRNA-flanked sgRNA induced higher level of RFP expression when compared to ribozyme-flanked sgRNAs (**Fig. 1b, c and Supplementary Fig. 1a, b, c**).

We then knocked in the tRNA-flanked sgRNA in the second intron of Lncenc1 in mouse embryonic stem cells (ESCs) in which CAGGS-dCas9-VPR and TRE-RFP have been transgenically integrated (**Fig. 1d**). Lncenc1 is a lncRNA specifically expressed in mouse ESCs^5^. In ESCs with GRIT successfully integrated (Lncenc1-GRIT ESCs), we observed high level of RFP expression (**Fig. 1e, f**). In addition, the knock-in of tRNA-sgRNA have little effect on the expression of Lncenc1 and pluripotency genes including Nanog, Oct4 (also known as Pou5f1) and Sox2 (**Supplementary Fig. 2a**). Importantly, the transcription activity of Lncenc1 locus was found not affected based on qPCR analysis of pre-mRNA of Lncenc1 (**Supplementary Fig. 2a**).

Lncenc1 is downregulated during ESC differentiation^5^. To check whether GRIT can report the expression of Lncenc1 during ESC differentiation, we induced differentiation of Lncenc1-GRIT ESCs with all-trans retinoid acids (ATRA) and measured RFP expression during differentiation process. Interestingly, RFP was significantly decreased upon ATRA induced differentiation (**Fig. 1g, h and Supplementary Fig. 2b**). More importantly, RFP level was highly correlated to the mRNA and pre-mRNA of Lncenc1 (Correlation coefficient for RFP versus Lncenc1 RNA~0.81, p value=0.0014). Furthermore, by comparing the expression of Lncenc1 and Oct4 during the differentiation process of wild type and Lncenc1-GRIT ESCs (**Fig. 1i and Supplementary Fig. 2c**), we concluded that knock-in of tRNA flanked sgRNA does not impact the differentiation potential of mouse ESCs. To check whether RFP level may reflect the stages of differentiation, we sorted out high and low RFP population in day 2 differentiated Lncenc1-GRIT cells and analyzed the expression of pluripotency genes (**Supplementary Fig. 3a**). Interestingly, Oct4, Nanog and Sox2 were indeed higher in RFP high cells than RFP low cells (**Supplementary Fig. 3b**). These results demonstrate the potential of GRIT to report the promoter activity of lncRNAs and as an indicator to monitor the differentiation status of stem cells.

NEAT1 is a lncRNA serving as a structural organizer of paraspeckle and has been shown to play important functions from gene regulation to cancer progression^6^. In addition, NEAT1 is an intronless gene. We constructed NEAT1-GRIT ESCs by inserting a mini-intron containing tRNA flanked sgRNA (**Fig. 1j**). As expected, RFP was significantly induced in NEAT1-GRIT ESCs (**Fig. 1k, l**). In addition, the insertion of mini-intron did not affect the expression of NEAT1 RNA and pluripotency genes including Oct4, Nanog, Sox2 and Klf4 (**Supplementary Fig. 4a**). We then performed embryoid body differentiation of NEAT1-GRIT ESCs and sorted out RFP high cells at day 8 (**Supplementary Fig. 4b**). As expected, qRT-PCR analysis showed that RFP high cells express higher level of NEAT1 (**Fig. 1m**). In addition, mesoderm marker T brachyurary was highly upregulated in RFP high cells (**Fig. 1m**), indicating that high NEAT1 expression may mark certain cell lineages from mesoderm.

In summary, we constructed an lncRNA reporter GRIT with CRISPR-on system by insertion of a tRNA-flanked sgRNA in endogenous lncRNA loci. We showed that GRIT is useful for tracking ESC differentiation and labelling specific cell lineages. A recent study from Gao *et al*. has reported a similar design which they named as Ents^7^. Ents uses Suntag-P65-HSF1 instead of VPR to activate gene expression and optimized mini-CMV-mCherry as a reporter. Different from traditional methods to report promoter activity by knocking in a protein such as GFP or luciferase, both Ents and GRIT strategies knock in a smaller DNA fragment that will not change the coding status of lncRNAs. In addition, by putting sgRNA in an intron, GRIT may have little impact on the expression of targeted lncRNAs, therefore not affecting the function of lncRNAs. This may be important for certain lncRNAs with essential functions. Finally, when sgRNAs are designed to target endogenous DNA locus, GRIT may be utilized to edit or manipulate the expression of endogenous genes. We expect that GRIT will be applicable in uncovering molecular mechanisms regulating the transcription of lncRNA, tracking cell fate switch during differentiation or disease progression and integrating the promoter activity of various RNAs for synthetic biology applications.

## Acknowledgments

We would like to thank Dr. Lu-Feng Hu and other members of Wang laboratory for discussion of the project. This study was supported by The National Key Research and Development Program of Ministry of Science and Technology of the People’s Republic of China [2016YFA0100701 and 2018YFA0107601] and National Natural Science Foundation of China [91940302, 31821091 and 32025007] to YW.

## Contributions

YTZ performed all the experiments. YTZ and YW interpretated the data. YW conceived and supervised the project and wrote the manuscript.

## Corresponding author

Correspondence to Yangming Wang.

## Supplementary Information

**Figure S1.**
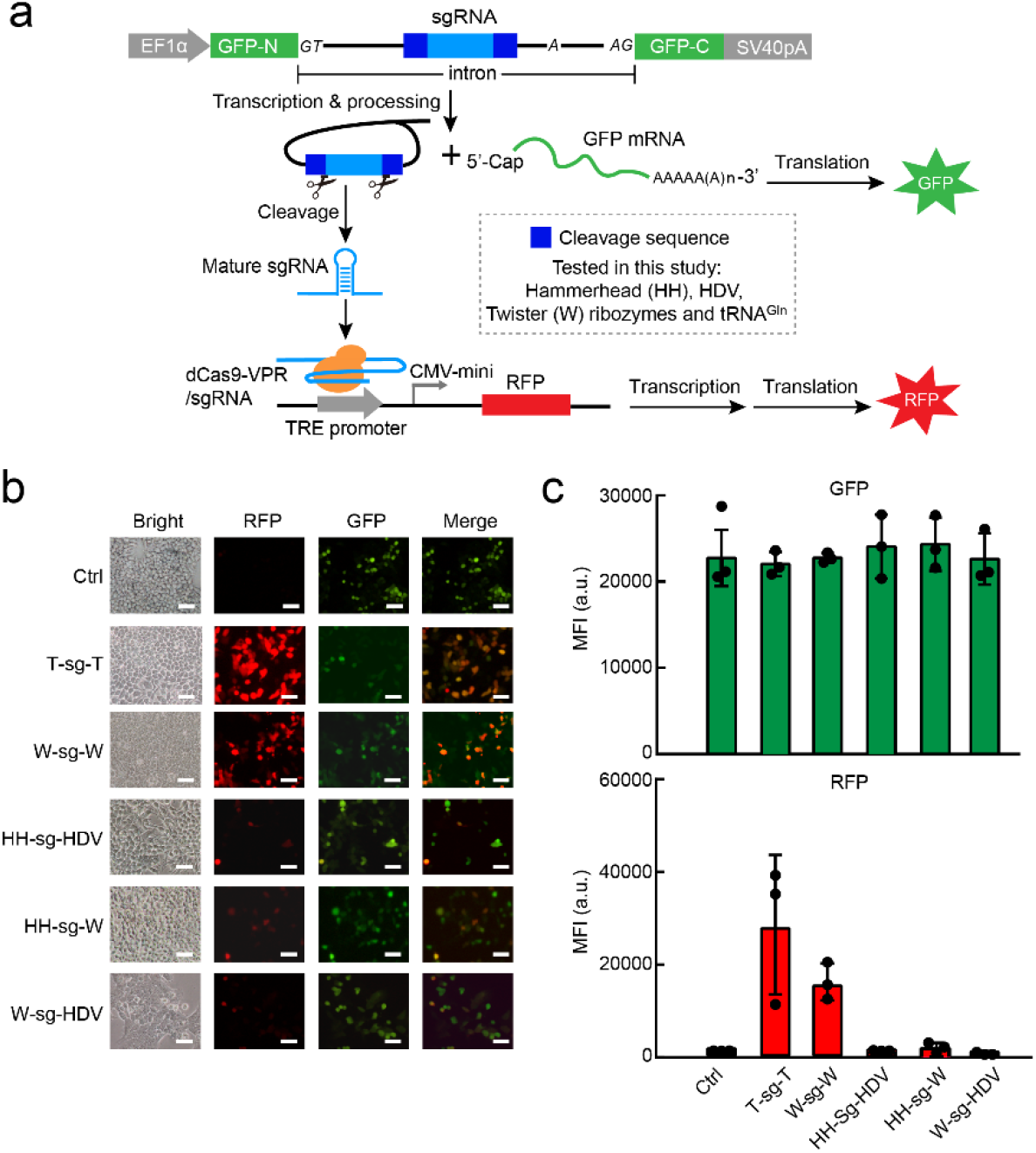
tRNA and twister ribozyme sequences flanking sgRNAs in GRIT lead to high RFP induction. **a)** Schematic representation of GRIT in an exogenous GFP gene. The pre-sgRNA in an intron cassette is inserted in GFP sequence driven by EF1alpha promoter. The intron was derived from the second intron of human *RPL18* gene. **b)** Representative microscopy images showing the expression of RFP and GFP for tested GRIT cassettes in HEK293T. Scale bar, 50μm. **c)** RFP and GFP intensity of GRIT cassettes overexpressed in HEK293T. Shown are mean ± SD, n = 3 independent experiments.

**Figure S2.**
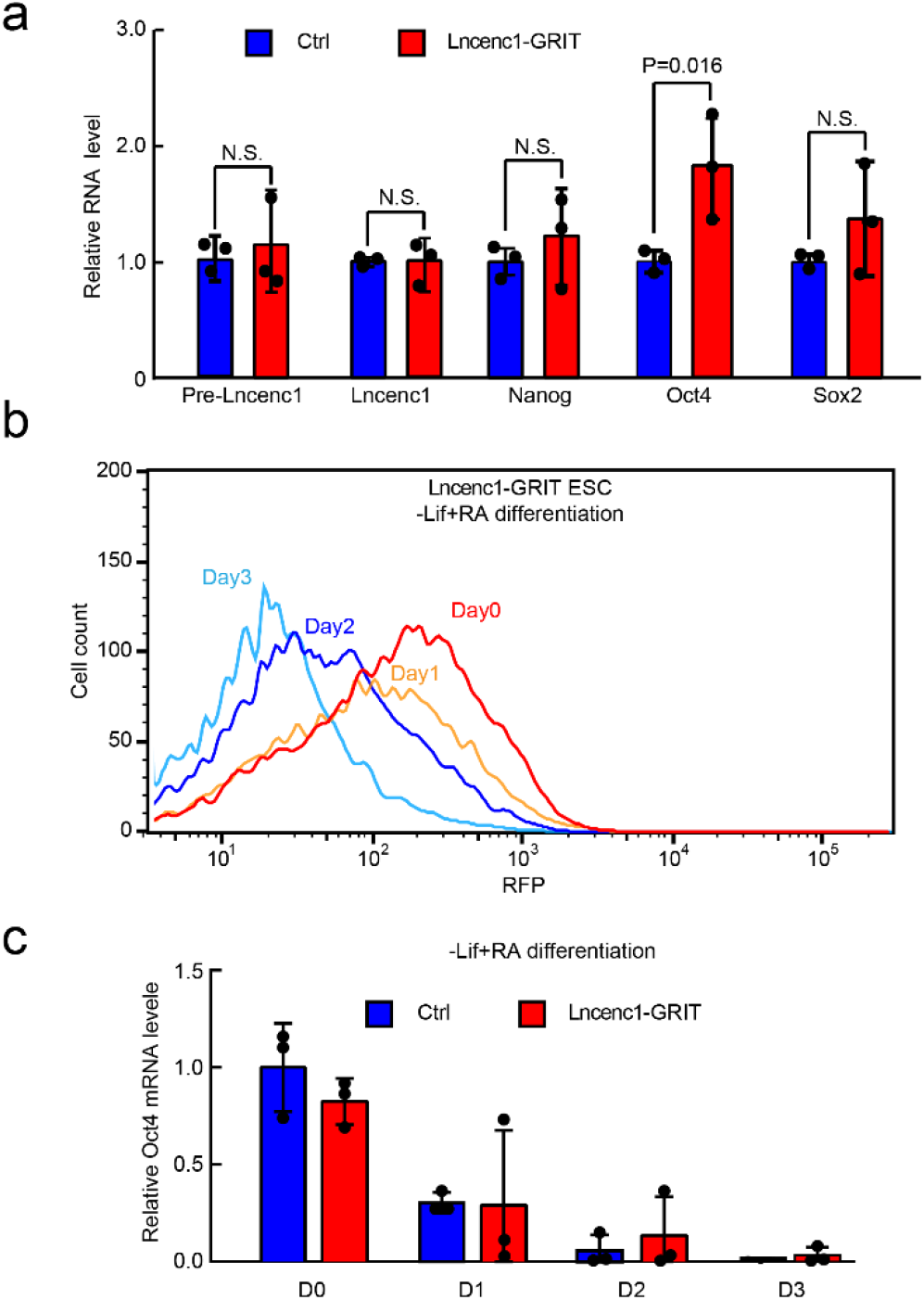
GRIT has little impact on the expression of Lncenc1 or the differentiation process of ESCs. **a)** RT-qPCR analysis of control and Lncenc1-GRIT ESCs. Control ESCs refer to embryonic stem cell line stably expressed dCas9-VPR and TRE3G-RFP with no GRIT knocked in. β-actin mRNA was used as a control. Data were normalized to control ESCs. Shown are mean ± SD, n = 3 independent experiments. The *p*-value was calculated by two-tailed Student’s *t* test. **b)** Representative flow cytometry analysis of undifferentiated and ATRA differentiated Lncenc1-GRIT cells. **c)** RT-qPCR analysis of Oct4 during differentiation process of control and Lncenc1-GRIT ESCs. β-actin mRNA was used as a control. Data were normalized to control ESCs. Shown are mean ± SD, n = 3 independent experiments.

**Figure S3.**
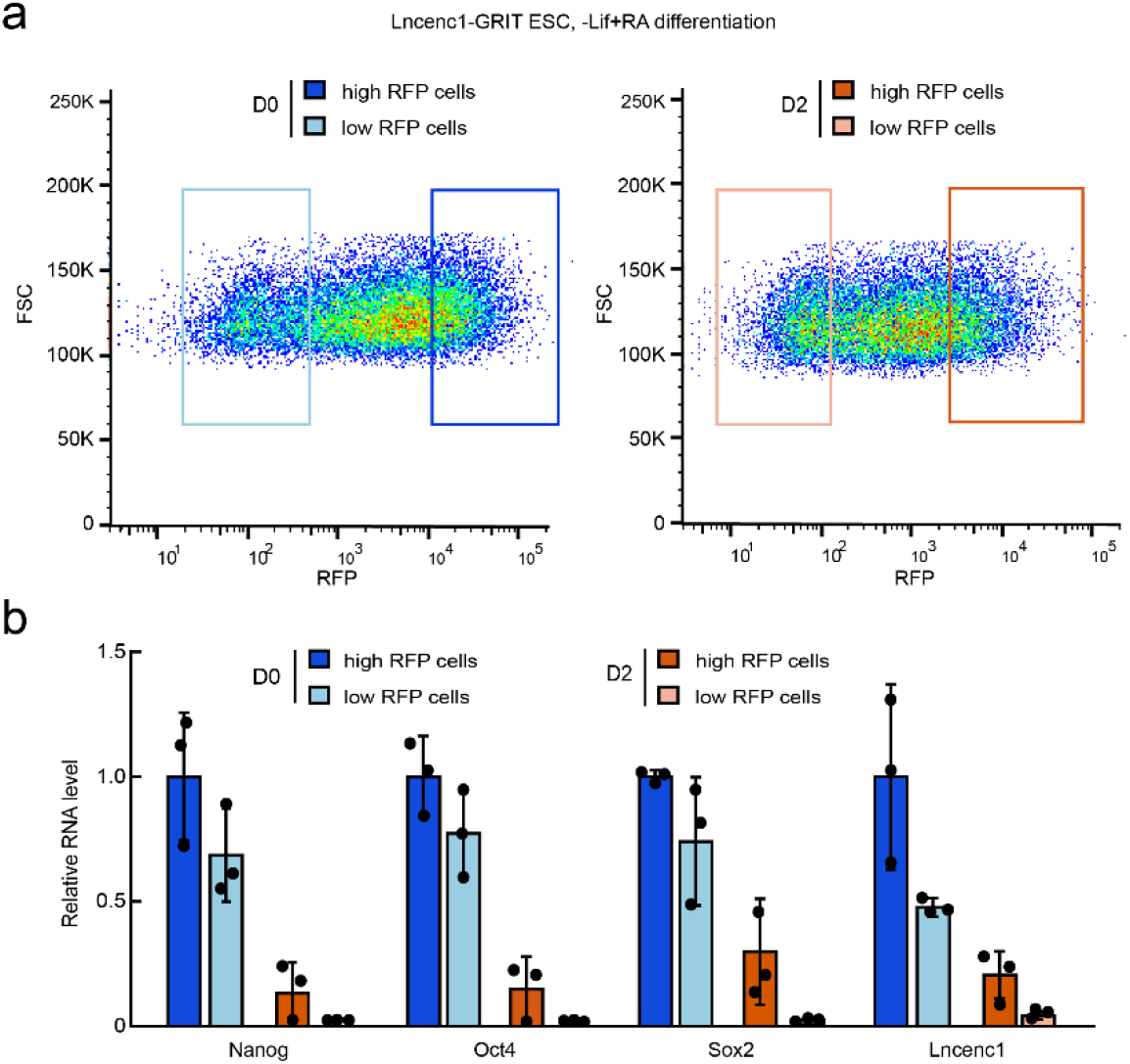
Lncenc1-GRIT can track the differentiation status of ESCs. **a)** Representative flow cytometry scatter plot and sorting gates of RFP high and low cells for undifferentiated and differentiated Lncenc1-GRIT ESCs. **b)** RT-qPCR analysis of RFP high and low undifferentiated and differentiated Lncenc1-GRIT ESCs. Data were normalized to β-actin gene and then undifferentiated high RFP intensity ESCs. Shown are mean ± SD, n = 3 independent experiments.

**Figure S4.**
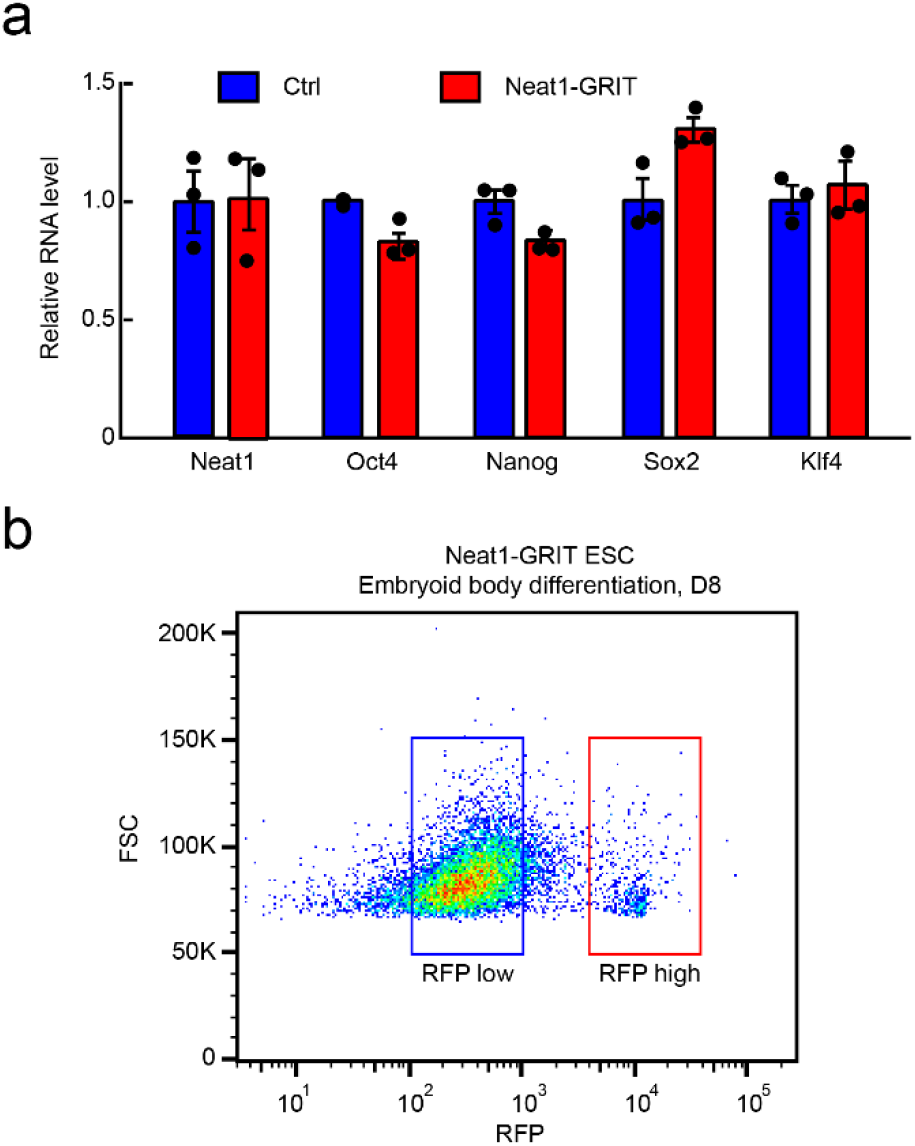
GRIT can track the expression of Neat1 in mouse ESCs. **a)** Representative flow cytometry scatter plot and sorting gates of RFP high and low cells from day 8 embryoid bodies of Neat1-GRIT ESCs. **b)** RT-qPCR analysis of control and Neat1-GRIT ESCs. Control ESCs refer to embryonic stem cell line stably expressed dCas9-VPR and TRE3G-RFP with no GRIT knocked in. β-actin mRNA was used as a control. Data were normalized to control ESCs. Shown are mean ± SD, n = 3 independent experiments.

## References

1 Djebali S, Davis CA, Merkel A et al. Landscape of transcription in human cells. Nature 2012; 489:101–108.

2 Kopp F, Mendell JT. Functional Classification and Experimental Dissection of Long Noncoding RNAs. Cell 2018; 172:393–407.

3 Wang XW, Hu LF, Hao J et al. A microRNA-inducible CRISPR-Cas9 platform serves as a microRNA sensor and cell-type-specific genome regulation tool. Nature cell biology 2019; 21:522–530.

4 He X, Wang Y, Yang F et al. Boosting activity of high-fidelity CRISPR/Cas9 variants using a tRNA(Gln)-processing system in human cells. The Journal of biological chemistry 2019; 294:9308–9315.

5 Sun Z, Zhu M, Lv P et al. The Long Noncoding RNA Lncenc1 Maintains Naive States of Mouse ESCs by Promoting the Glycolysis Pathway. Stem cell reports 2018; 11:741–755.

6 Wang Y, Chen LL. Organization and function of paraspeckles. Essays in biochemistry 2020; 64:875–882.

7 Gao N, Hu J, He B et al. Endogenous promoter-driven sgRNA for monitoring the expression of low-abundance transcripts and lncRNAs. Nature cell biology 2021.

8 Yoshioka S, Fujii W, Ogawa T, Sugiura K, Naito K. Development of a mono-promoter-driven CRISPR/Cas9 system in mammalian cells. Scientific reports 2015; 5:18341.

9 Litke JL, Jaffrey SR. Highly efficient expression of circular RNA aptamers in cells using autocatalytic transcripts. Nature biotechnology 2019; 37:667–675.

